# Skin Lesions Classification Using Deep Learning Based on Dilated Convolution

**DOI:** 10.1101/860700

**Authors:** Md. Aminur Rab Ratul, M. Hamed Mozaffari, Won-Sook Lee, Enea Parimbelli

**Affiliations:** School of Electrical Engineering and Computer Science, University of Ottawa, Ottawa, Canada; Telfer School of Management, University of Ottawa, Ottawa, Canada

**Keywords:** Dilated Convolution, Medical Image Analysis, Skin Lesion Classification, Deep Learning, Convolutional Neural Network

## Abstract

The prediction of skin lesions is a challenging task even for experienced dermatologists due to a little contrast between surrounding skin and lesions, the visual resemblance between skin lesions, fuddled lesion border, etc. An automated computer-aided detection system with given images can help clinicians to prognosis malignant skin lesions at the earliest time. Recent progress in deep learning includes dilated convolution known to have improved accuracy with the same amount of computational complexities compared to traditional CNN. To implement dilated convolution, we choose the transfer learning with four popular architectures: VGG16, VGG19, MobileNet, and InceptionV3. The HAM10000 dataset was utilized for training, validating, and testing, which contains a total of 10015 dermoscopic images of seven skin lesion classes with huge class imbalances. The top-1 accuracy achieved on dilated versions of VGG16, VGG19, MobileNet, and InceptionV3 is 87.42%, 85.02%, 88.22%, and 89.81%, respectively. Dilated InceptionV3 exhibited the highest classification accuracy, recall, precision, and f-1 score and dilated MobileNet also has high classification accuracy while having the lightest computational complexities. Dilated InceptionV3 achieved better overall and per-class accuracy than any known methods on skin lesions classification to the best of our knowledge while experimenting with a complex open-source dataset with class imbalances.

## I. Introduction

In recent times, skin cancer is one of the most common forms of cancer not only in the USA (5 million cases per annum) but also in all over the world [1-5]. Squamous cell carcinoma, melanoma, intraepithelial carcinoma, basal cell carcinoma are some of the usual kinds of skin lesions [6-8], but among them, melanoma is most dangerous and extremely cancerous (over 9000 deaths in 2017 only in the USA) [3]. Early diagnosis of melanoma can cure nearly 95% cases [9], and through dermoscopy, the accuracy of skin lesions treatment will be 75%-84% [10-12].

The manual skin lesions detection system is human-labor intensive, which needs magnifying and illuminated skin images to improve the clarity of spots [10, 13]. ABCD-rule (Asymmetry, Border, Irregularity, Color variation, and Diameter), 3-point checklist, 7-point checklist, and Menzies method are several procedural algorithms to boost the dermoscopy and observe the malignant melanoma in the very early stage [11, 14] however, and many clinicians steadily rely on their experiences [15]. The manual dermoscopy imaging procedure is more prone to mistake because it needs years of experience over difficult situations, vast amounts of visual exploration, similarities, and dissimilarities between different skin lesions.

In recent times, deep learning (begin with AlexNet [16] in 2012) provides many computerized automated systems to detect, classify, and diagnosis of several diseases through medical image analysis [17]. Last few years, dermoscopy produce a significant amount of well-annotated skin lesions images that help supervised machine learning techniques actively to classify, predict, and detect different skin wound [10, 18-20]. Hence, deep learning-based medical image analysis tools can be useful to assist the dermatologist to emphasis on several areas like skin lesion segmentation, classification, and detection.

Here, we proposed a deep learning method, namely dilated or, atrous convolution, with transfer learning to classify seven different class skin lesions. Compared to traditional CNN, we used dilated convolution to increasing accuracy with the same computational complexities. We choose four pre-trained deep learning architectures such as VGG16, VGG19, MobileNet, and InceptionV3 and select a different strategy to put different dilation rates in separate layers. To the best of our knowledge, we are the first who proposed the approach to employ different dilation rates in the different layers of InceptionV3 and MobileNet network and achieve better overall performance than the original architectures. Moreover, we utilize a fine-tuning technique to train these proposed architectures. We use the HAM10000 dataset to train, validation, and test, which contains 10015 dermatoscopic images of seven skin lesions like Vascular lesions, Actinic Keratoses, Benign keratosis-like lesions, Dermatofibroma, melanoma, melanocytic nevi, and Basal cell carcinoma [21]. Melanoma is extremely dangerous, Basal cell carcinoma, Actinic keratoses can be cancerous, and the other skin lesions in this dataset are benign.

The rest of the paper ordered as follows: in section 2, discussion on related work. Data utilization and Methodology and illustrated in section 3 and section 4 respectively. The performance analysis of the models presented in section 5. Finally, section 6 comprises the conclusion and the future work part.

## II. Related work

In previous times, several types of research had been done to detect skin cancer. Those mainly based on splitting and merging region, clustering, supervised learning, and thresholding, but every work have many pros and cons [22-24]. Celebi et al. [25] proposed four ensemble thresholding methods to skin lesion border detection, and Sigurdsson et al. [26] developed a probabilistic feature extraction model with a feedforward neural network to classify skin lesions. Barata et al. constructed two approaches, namely global features and local features (utilize BoF (Bag of Features)), to detect melanoma using dermoscopy images [27]. To classify skin lessons, She et al. [28] use several features like diameter, color, border, and asymmetry. To attain skin lesion segmentation, several pre-processing methods such as artificial removal, color transformation, lesion localization, contrast enhancement can be employed [29].

Recently several deep learning models have been build for classification, detection, and segmentation. Hekler et al. [30] execute deep learning methods to classify histopathologic diagnosis of melanoma, to augment the human evaluation, and contrasted the outcome with skillful histopathologists. Esteva et al. [31] implemented pre-trained inceptionv3 for nine class classification where they used a labeled dataset by dermatologists, which have 3374 dermoscopy images, 129,450 clinical images, and achieve 72.1 ± 0.9% (mean ± s.d.) accuracy. Harangi et al. [32] utilize an ensemble DCNN (deep convolutional neural network) method, where they combine the result of four different architectures with improving the accuracy of the ISBI 2017 [1] dataset. Rather than training CNN (Convolutional Neural Network) from scratch, Kawahara et al. [33] attempted to employ pre-trained ConvNet as their feature extractor, and they classify ten classes of non-dermoscopic skin images. Xie and Bovik [34] displayed a skin lesion segmentation method where a self-generating CNN merged with a genetic algorithm. Recently, Carcagni et al. [35] proposed a research work based on a multilevel DensNet network to classify seven different skin lesions of HAM10000. Gomez et al. [36] published a skin lesion segmentation architecture, namely Independent Histogram Pursuit (IHP), where they tested their method on five different dermatological datasets and obtained 97% precision on segmentation. Yunfei et al. [37] proposed a deep residual model for classification of skin pigmented lesions, which upgraded by class weight update dynamically, Excitation, and Squeeze module in batches. A new fully CNN segmentation method proposed with new pooling layers for skin lesions region in [38]. In [39], Mask R-CNN and U-net utilized for skin lesion segmentation in dermoscopic images, particularly on ISIC 2017 dataset. Yu et al. [40] to differentiate melanoma images from non-melanoma, a very deep residual CNN has proposed.

In the present time, many classification deep learning models proposed for dermoscopy images; however, to build a future-oriented model, we can make improvements in several different areas. So, for this work, we implement a hugely popular method called dilated convolution for classifying seven skin lesions with transfer learning techniques. Previously, dilated convolution utilized in instance segmentation [43], audio generation [46], optical flow [42], question answering [41], and object detection [44,45].

## III. Dataset

Here, we use HAM10000 dataset [1, 22] which contains 10015 dermatoscopic skin lesion images of seven classes (Melanocytic nevi (6705 images), Melanoma (1113 images), Benign keratosis-like lesions (1099 images), Basal cell carcinoma (514 images), Actinic keratoses (327 images), Vascular lesions (142 images), and Dermatofibroma (115 images)). We applied the stratified method to split HAM10000 into training (80% or, 8011), validation (10% or, 1002), and test (10% or, 1002) sets so that we can maintain the ration between every class because this is a class imbalanced dataset. In table 1, we manifest the training, validation, and test sets after the stratified technique.

**TABLE I.** Training, validation, and testing sets after employ the stratified technique on HAM10000

| Class Name | Training (80%) | Validation (10%) | Testing (10%) |
| --- | --- | --- | --- |
| Actinic keratoses | 261 | 33 | 33 |
| Basal cell carcinoma | 412 | 51 | 51 |
| Benign keratosis-like lesions | 879 | 110 | 110 |
| Dermatofibroma | 91 | 12 | 12 |
| Melanocytic Nevi | 5363 | 671 | 671 |
| Vascular lesions | 114 | 14 | 14 |
| Melanoma | 891 | 111 | 111 |
| Total Images | 8011 | 1002 | 1002 |

### A. Data Pre-Processing

To lessen the computational cost of our proposed architecture, we resize and rescale the image size from 600 × 450. To maintain the same contorted, at first, we crop the center part and then reduce the dimension of the images by the following width to height ratio. We inspected different combinations of image shapes (320 × 320,512 × 384, 256 × 192, 128 × 96, and 64 × 64) for VGG16, VGG19, and InceptionV3. MobileNet only takes four forms of static square images ((128 × 128), (160 ×160), (192 × 192), (224 × 224)) [47]. Therefore, based on time complexity, space complexity, and accuracy, we select image dimensions for the first three models and only for MobileNet. Finally, to normalize our data, we divide each pixel of image values by 255.0 to rescale pixel values into the 0-1 range.

### B. Data Augmentation

Deep learning models require a decent amount of data to produce good results [48], so we use different data augmentation processes to train our model. We train our models for 200 iterations and able to create new transformed 1,602,200 (220 × 8011) images for the whole training process. Hence, for each iteration, 8011 newly transformed augmented images have been provided. Additionally, for the validation set, we employ the same strategy and able to produce 1002 new augmented images to validate our training. In our training set, we applied many geometrical transformations for data augmentation. We employ vertical and horizontal flipping with a probability of 0.50 to transform the images. To handle the off-centered objects, random width and height shifting with range 0.20 have utilized. Then, the zoom range (−0.2, +0.2) was used to randomly zooming in inside. Lastly, we applied shear mapping to supplant the image horizontally (*x* + 0.2,*y*) or vertically (*x,y* + 0.2).

## IV. Methodology

Four popular deep learning models, namely VGG16 [49], VGG19, MobileNet [47], and InceptionV3 [50] with atrous or dilated convolution instead of traditional convolution, had been used to build an accurate automated model for the dermatologist. To strengthen the accuracy, we utilized the transfer learning technique with a pre-trained ImageNet dataset for these dilated CNN networks.

### A. Dilated Convolution

Initially, researchers invented an algorithm, namely “hole algorithm” or “algorithme à tours” for wavelet transformation [51], but right now in the deep learning area is known as “atrous convolutional” or, “Dilated Convolution”.

The dilated convolution expands the kernel’s field of view with the same computational complexities by insert “hole” or zeros between the kernel of each convolutional layer. Therefore, it can use for those applications which cannot bear bigger kernels or, many convolutions, however, require a wide field of vision.

The scenario of dilated convolution in the 1D field:

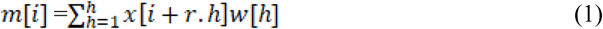

Here, for every location of *i*, the output is *y*. Furthermore, *x[i]* is the input signal where *x* also referred to as a feature map. Besides, *w[h]* filtered with the length of *h*, and *r* is corresponding to the dilation rate with which we sample input signal *x[i]*. In the standard convolution r =1 but the dilated convolution rate of r is always bigger than 1. An intuitive and easy way to comprehend dilated convolution is that push (r-1) zeros between every two consecutive filters in the standard convolution. In a standard convolution, the kernel or filter size is *n* × *n*, then the resulting dilated convolution, the filter or kernel size will be *n*_*d*_ × *n*_*d*_ where *n*_*d*_ *= n + (n-1). (r-1)*.

One of the main reasons to implement this method is without missing any coverage or resolution, dilated convolutions support exponentially enlarging receptive fields [52].

### B. Dilated VGG16 and Dilated VGG19 Architecture

Both VGG16 and VGG19 have five blocks of convolutional layers were with an equal number of parameters to expand the context view of filters where we modify the dilation rate of these layers. Output feature map will shrink (output stride increasing) for any standard convolution and pooling if we go deeper in any model, which is harmful for classification because, in the deep layers, spatial information will be missing. With dilated convolution without increase computational complexities, we can achieve a larger output feature map, which is proved to be appropriate for skin lesion classification in terms of accuracy.

Both the model has 3 × 3 kernel size in every layer, and without increasing kernel size, we can enhance the receptive field dimension by adding different dilation rates in the existing layer in our proposed architectures. The input layer size is 192 × 256 × 3 (*height of the image* × *width of the image* × *RGB*).

The initial block of these networks have *dilation rate* = 1, then from block two to five have dilation rate 2, 4, 8, and 16 respectively. To implement this technique, we mostly inspired by different multi-grid models where we find a hierarchy of several different sizes of grid [53-56], many semantic image segmentation models [57, 58], in [58] Chen et al. pick distinct dilation rates within block4 to block7 in the proposed ResNet model. Seemingly, we utilize VGG networks to adopt different dilation rates in different blocks.

In the all convolutional layer, we used rectified linear units (ReLUs) as activation function and max-pooling used for downsampling in between every convolutional block. After the last convolutional and max-pooling layer, we run *global max-pooling* operation which takes tensor with shape *h* × *w* × *d* (*h* ×*w*= spatial dimensions, *d = number of feature maps*) and provides output tensor with shape. Then, we add two fully connected layers in these models with 512, 7 (dataset has seven classes) filters respectively with a dropout layer () in between which utilize as a regularizer function to substantially weaken the overfitting rate and computationally reasonable at the same time [59]. “RELU” is the activation function for the first dense layer, and the last one has “SoftMax”. From Figures 3 and 4, we can notice the details visualization of dilated VGG16 and VGG19.

**Fig. 1.**
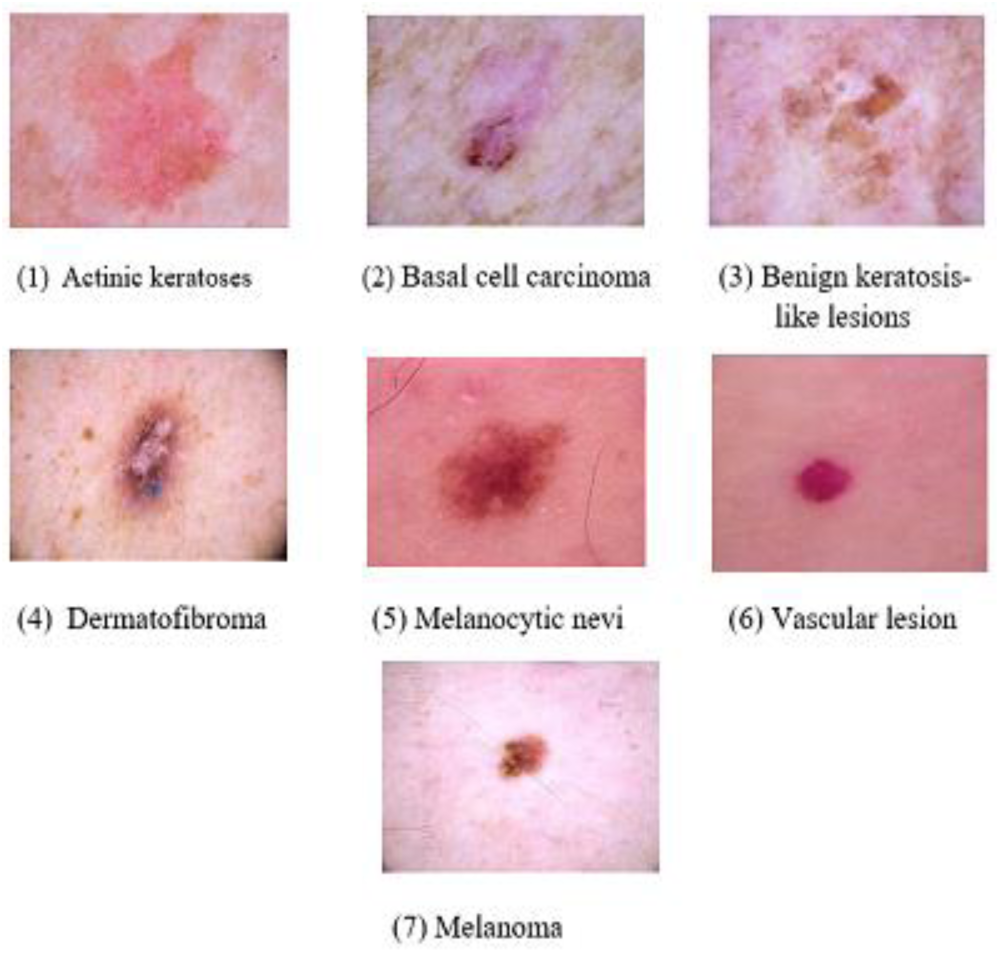
Seven different Skin lesions from HAM10000 dataset: Actinic keratoses, Basal cell carcinoma, Benign keratosis-like lesions, Dermatofibroma, Melanocytic Nevi, Vascular lesions, Melanoma

**Fig. 2.**
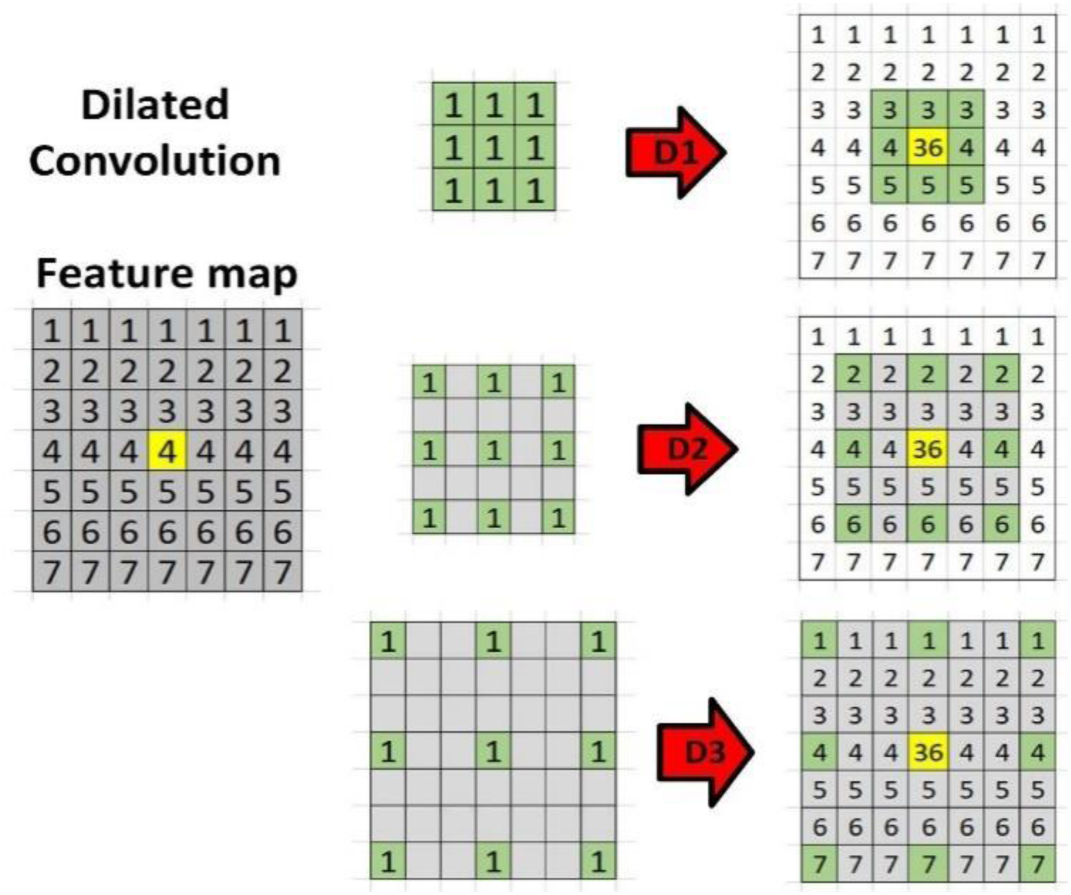
A scenario of dilated convolution with kernel size 3×3. From the top: (a) the situation of the standard convolutional layer, and the dilation rate is (1,1). (b) when the dilation rate becomes (2,2), and the receptive field enhances. (c) in the final case, the dilation rate is (3,3), and the receptive field extends more than the situation b

**Fig. 3.**
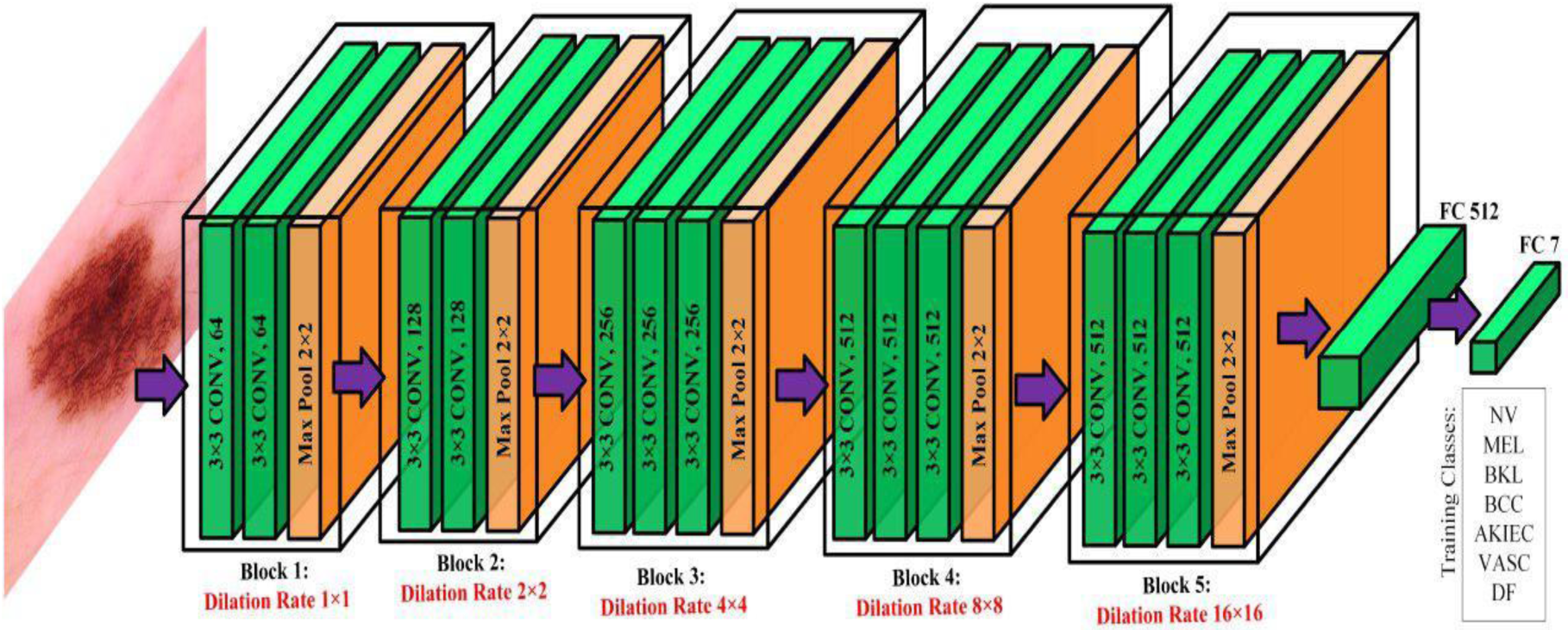
Dilated VGG16 model with different dilation rate on every block

**Fig. 4.**
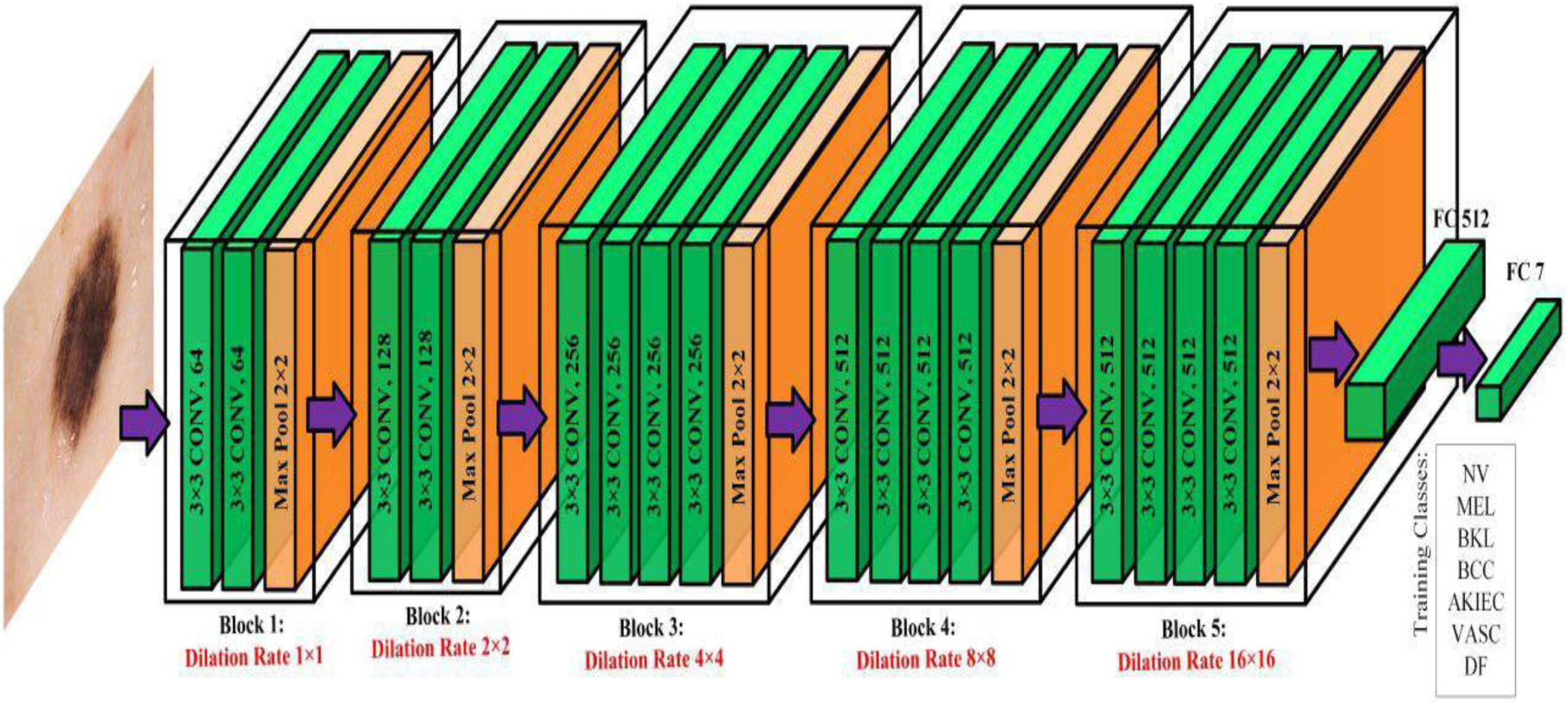
Dilated VGG19 model with different dilation rate on every block

### C. Dilated MobileNet Architecture

MobileNet was constructed to provide small, very low latency, and computationally sound model for embedded mobile vision applications [47]. MobileNet has three kinds of convolutional operation: standard convolution, pointwise convolution, and depthwise convolution. We take five depthwise layers to implement the dilated convolution, and every layer has a stride rate (2,2). Among these depthwise layers, the first two layers have a dilation rate (1,1); however, for the third and fourth layers, we placed a dilation rate (2,2). Furthermore, for the final depthwise layer, we concatenate three depthwise 2D convolution layers parallelly with a dilation rate of 4,8,16, respectively. Finally, we concatenate these three 2D layers and produce the fifth depthwise convolutional 2D layer. Originally, every depthwise layer of MobileNet has but implementing different dilation rates in distinct depthwise layers of MobileNet architecture is new, and we first propose this approach.

After all the convolution operation (standard, depthwise, and pointwise), from the last pointwise layer, we take the feature map and employ the *global average pooling* (GAP) method. *Global average pooling* converts the feature map size into 1 × 1 × *d* from *h* × *w* × *d*, and here this method takes average value from the spatial dimension of the feature map (*h* × *w = spatial dimension*). GAP has several advantages; such as elude overfitting in the layer, in the input feature map, it exhibits more robust characteristics to the spatial translations [60]. The classifier part of the fully connected part is the same as the VGG networks. There are two fully connected layers, and in between, there is a dropout layer.

### D. Dilated InceptionV3

GoogleNet [62] produces InceptionV3 architecture to perform efficiently under the strict limitations of memory and computation. GoogleNet [62] first displayed back in 2014 in ImageNet [61] competition.

InceptionV3 has three main parts in its networks, like factorizing convolution, auxiliary classifier, and coherent grid size reduction. Again we can divide factorizing convolution into three sections. First, instead of using a big convolution, factorizing it into the two smaller convolutions is computationally inexpensive. For instance, in InceptionV3 two 3 × 3 convolutions used in the place of one 5 × 5 convolution because two 3 × 3 filters would cost us 18 (3 * 3 + 3 * 3), whereas only one 5 × 5 filter cost us 25 (5 * 5). So, for this model, we named this section as Module A and inserted *dilation rate* = 2 on each convolutional layer of this module. Secondly, in the place of *m* × *m* filter shape (3 * 3 = 9), this network using 1 × *m* + *m* × 1 convolutions combination (1 * 3 + 3 * 1 = 6), which is 33% cheaper in terms of parameters. We call this section Module B and put *dilation rate* = 2 in every layer of this module. Finally, there is a section present in InceptionV3 to present the high dimensionality, and we called it Module C and placed *dilation rate* = 2 in the convolutional layer. To prevent the loss of important information, filter banks have been enlarged in this part where filter banks become wider rather than go deeper. Original InceptionV3 has *dilation rate* = 1 in every module. To the extent of our knowledge, we are the first who suggested putting *dilation rate* = 2 in different modules and finally achieve a better outcome than the original InceptionV3.

Lastly, after the last module of this model, we use *global max pooling*, and the classifier or fully connected part is the same as our previous implementation. From figure 3, we can find the details process of dilated InceptionV3.

### E. Transfer Learning and Fine Tune Technique

We apply a similar fine-tuning procedure for all the dilated networks which are already pre-trained with ImageNet dataset [61] so that every layer of these models has some weight from ImageNet before initiate our training. Firstly, we detach fully connected layers or top layers part of the existing models. Next, we attach a new fully-connected part as a classifier (this part we already described). To perform feature extraction, we freeze all layers instead of the classifier part and run this fully connected part for five epochs to give some weight; otherwise, the gradient would be so high because all the freeze layers have some pre-trained weight from ImageaNet. Finally, we unfroze the other layers and run the training for 200 epochs to put the extra weight of the HAM10000 dataset on every layer of our proposed models.

## V. Result

To construct our models, mainly we use Keras for the frontend development and TensorFlow for the backend. Next, Pandas and Scikit Learn utilized respectively for data pre-processing, and to evaluate these proposed models. Training every model for 200 iterations and take 32 as the mini-batch size. The models executed on Intel Core i7-8750H with 4.1 GHz and an NVIDIA GeForce GTX 1050Ti GPU. Adam ^64^ optimizer used as the optimization function with a learning rate 10^−4^ initially. One callback function utilized to lessen the learning rate factor by (0.1)^.5^ during the training when the loss of validation is not diminishing for seven iterations. Thus, the new learning rate:

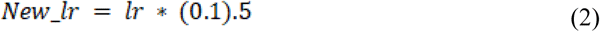

*New_lr* = new learning rate; *lr* = present learning rate. Here, the lower bound for the overall learning rate is 0.5*e* − 6.

### A. Top-1 Test Accuracy of Four Models

Before proposed these models, we tried numerous combinations for these four architectures. After the experiment with several combinations, we able to fixed which design for each model produces the best top1-accuracy and per-class accuracy. Furthermore, we examine different image resolution (64×64,128×96, 256×192, 320×320) for proposed VGG16, VGG19, and InceptionV3. Overall, these three networks produce the best outcome for 256×192 image resolution. On the other hand, among the different image size combinations for dilated MobileNet (128×128, 160×160, 192×192, 224×224) and 224×224 image shape provide the highest accuracy. From table 2, we can see the dilated InceptionV3 showed the foremost top-1 accuracy among these four models, and it displayed superior computational complexities. However, dilated MobileNet provides lightest computational complexities with only 3.7 million parameters.

**TABLE II.**
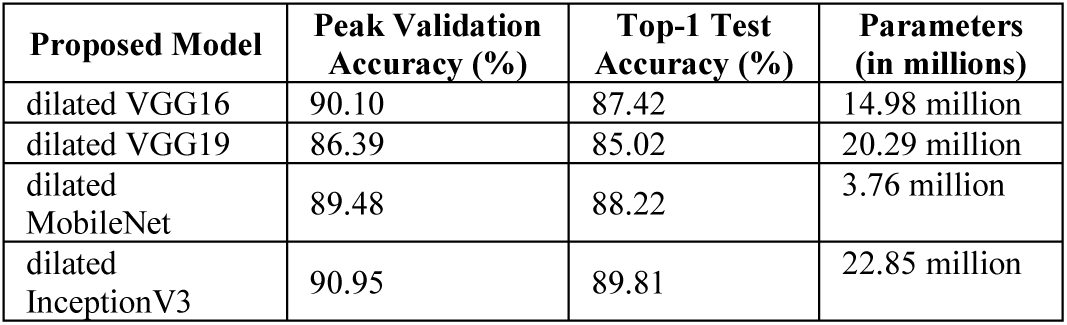
Comparison between four different dilated deep learning architectures based on peak validation accuracy, top-1 accuracy, and parameters

### B. Recall, Precision, and F-1 Score

HAM10000 is a high-class imbalanced dataset. So, evaluate the proposed architectures we displayed recall, precision, and F1 score for each model.

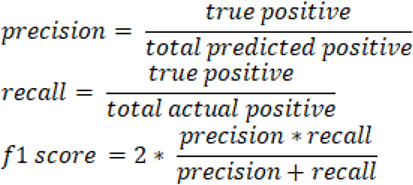

Precision means the percentage of the pertinent outcomes (how many picked entries are pertinent?). Recall, or sensitivity implies the rate of total apposite results accurately classified by the particular selected model (how many pertinent components are chosen?). F1-score means the harmonic mean of precision and recall. In table 3, we showed the recall, precision, f1-score of four proposed architecture. Furthermore, we displayed macro avg., micro avg., and weighted avg. of these evaluation parameters.

**TABLE III.** Precision, recall, F1-score, micro average, macro average, and weighted average. for the test set of the HAM10000 of proposed dilated VGG16

| Class Name | Precision | Recall | F1-Score |
| --- | --- | --- | --- |
| Actinic keratoses | 0.64 | 0.42 | 0.51 |
| Basal cell carcinoma | 0.80 | 0.84 | 0.82 |
| Benign keratosis like lesions | 0.72 | 0.76 | 0.74 |
| Dermatofibroma | 0.67 | 0.67 | 0.67 |
| Melanocytic Nevi | 0.94 | 0.96 | 0.95 |
| Vascular lesions | 1.00 | 0.79 | 0.88 |
| Melanoma | 0.72 | 0.66 | 0.69 |
| Micro Avg | 0.87 | 0.87 | 0.87 |
| Macro Avg | 0.78 | 0.73 | 0.75 |
| Weighted Avg | 0.87 | 0.87 | 0.87 |

**TABLE IV.** Precision, recall, F1-score, micro average, macro average, and weighted average. for the test set of the HAM10000 of proposed dilated VGG19

| Class Name | Precision | Recall | F1-Score |
| --- | --- | --- | --- |
| Actinic keratoses | 0.61 | 0.58 | 0.59 |
| Basal cell carcinoma | 0.79 | 0.65 | 0.71 |
| Benign keratosis like lesions | 0.66 | 0.82 | 0.73 |
| Dermatofibroma | 0.67 | 0.33 | 0.44 |
| Melanocytic Nevi | 0.93 | 0.93 | 0.93 |
| Vascular lesions | 0.91 | 0.71 | 0.80 |
| Melanoma | 0.67 | 0.65 | 0.66 |
| Micro Avg | 0.85 | 0.85 | 0.85 |
| Macro Avg | 0.75 | 0.67 | 0.70 |
| Weighted Avg | 0.85 | 0.85 | 0.85 |

**TABLE V.** Precision, recall, F1-score, micro average, macro average, and weighted average. for the test set of the HAM10000 of proposed dilated Mobilenet

| Class Name | Precision | Recall | F1-Score |
| --- | --- | --- | --- |
| Actinic keratoses | 0.79 | 0.45 | 0.58 |
| Basal cell carcinoma | 0.94 | 0.65 | 0.77 |
| Benign keratosis like lesions | 0.65 | 0.86 | 0.74 |
| Dermatofibroma | 0.90 | 0.75 | 0.82 |
| Melanocytic Nevi | 0.96 | 0.94 | 0.95 |
| Vascular lesions | 0.92 | 0.79 | 0.85 |
| Melanoma | 0.73 | 0.79 | 0.76 |
| Micro Avg | 0.88 | 0.88 | 0.88 |
| Macro Avg | 0.84 | 0.75 | 0.78 |
| Weighted Avg | 0.89 | 0.88 | 0.88 |

**TABLE VI.** Precision, recall, F1-score, micro average, macro average, and weighted average. for the test set of the HAM10000 of proposed dilated Inceptionv3

| Class Name | Precision | Recall | F1-Score |
| --- | --- | --- | --- |
| Actinic keratoses | 1.00 | 0.61 | 0.75 |
| Basal cell carcinoma | 0.90 | 0.71 | 0.79 |
| Benign keratosis like lesions | 0.71 | 0.84 | 0.77 |
| Dermatofibroma | 0.90 | 0.75 | 0.82 |
| Melanocytic Nevi | 0.93 | 0.97 | 0.95 |
| Vascular lesions | 0.91 | 0.71 | 0.80 |
| Melanoma | 0.83 | 0.70 | 0.76 |
| Micro Avg | 0.89 | 0.89 | 0.89 |
| Macro Avg | 0.88 | 0.75 | 0.81 |
| Weighted Avg | 0.89 | 0.89 | 0.89 |

**TABLE VII.** The comparison result of our proposed architectures with some existing models

| Source | Model | Avg. Recall | Avg. Precision | Avg. F-1 Score |
| --- | --- | --- | --- | --- |
| [40] | Original Densenet-121 | 0.49 | 0.36 | 0.30 |
| [40] | Combining Multilevel Learnings in a DenseNet (Ensemble DenseNet Architecture) | 0.88 | 0.76 | 0.82 |
| [65] | Fine Tune MobileNet | **0.89** | 0.83 | 0.83 |
| **Our Proposed Models** | Dilated VGG16 | 0.87 | 0.87 | 0.87 |
|  | Dilated VGG19 | 0.85 | 0.85 | 0.85 |
|  | Dilated MobileNet | 0.89 | 0.88 | 0.88 |
|  | Dilated InceptionV3 | **0.89** | **0.89** | **0.89** |

### C. Confusion Matrix

Because of the class imbalance issue, we need a confusion matrix to get a clear idea of our proposed models. Through this, we can able to evaluate where our models can make mistakes and confusion matrix used to sketch the performance of this architecture. For instance, in the second row of the confusion matrix of dilated VGG16 model, we find that it correctly labeled 43 images as Basal cell carcinoma; however, the architecture mistakenly labeled 2 images as Actinic Keratoses, 1 image as Benign Keratosis like lesions, 1 image as Dermatofibroma, 2 images as Melanocytic Nevi, and 5 images as Melanoma skin lesion. In figure 7, 8, 9, and 10 we provide four confusion matrix for four proposed models. In these four figures: AK= Actinic keratoses, BCC=Basal cell carcinoma, BK=Benign keratosis-like lesions, Der=Dermatofibroma, MN=Melanocytic Nevi, VL= Vascular lesions, Mel=Melanoma.

**Fig. 5.**
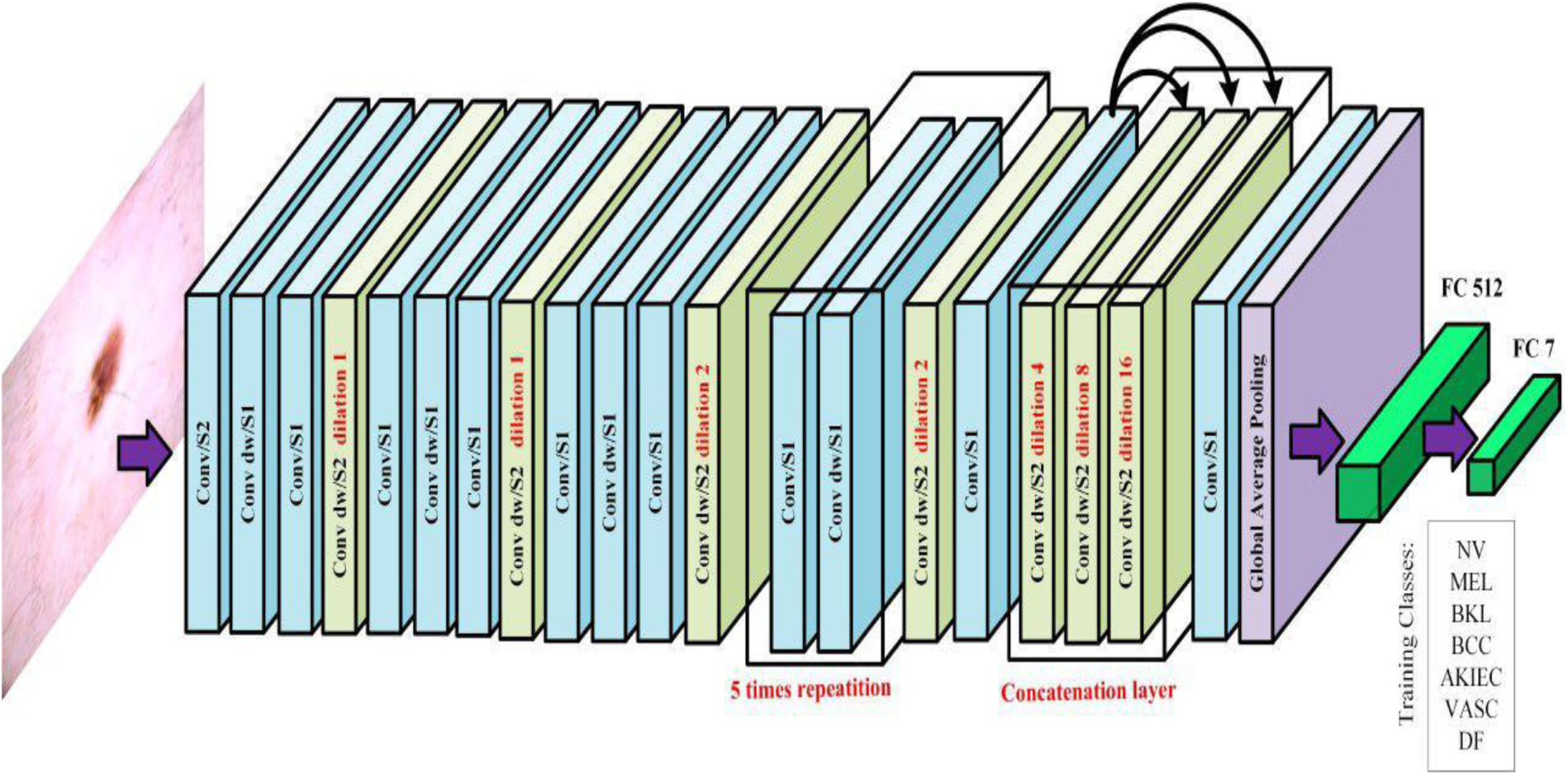
Dilated MobileNet architecture with different dilation rates on its depthwise convolutional layer, which contains stride (2,2). In the first two depthwise convolutional layers has a dilation rate (1,1). The third and fourth Depthwise layers experience dilation rate (2,2). For the final depthwise layer, we merge three parallel depthwise layers with dilation rate (4,4), (8,8), (16,16) respectively, and run concatenation operation to convert these three layers into one depthwise layer. In this figure, dw = depthwise; S1= stride (1,1); S2 = Stride (2,2)

**Fig. 6.**
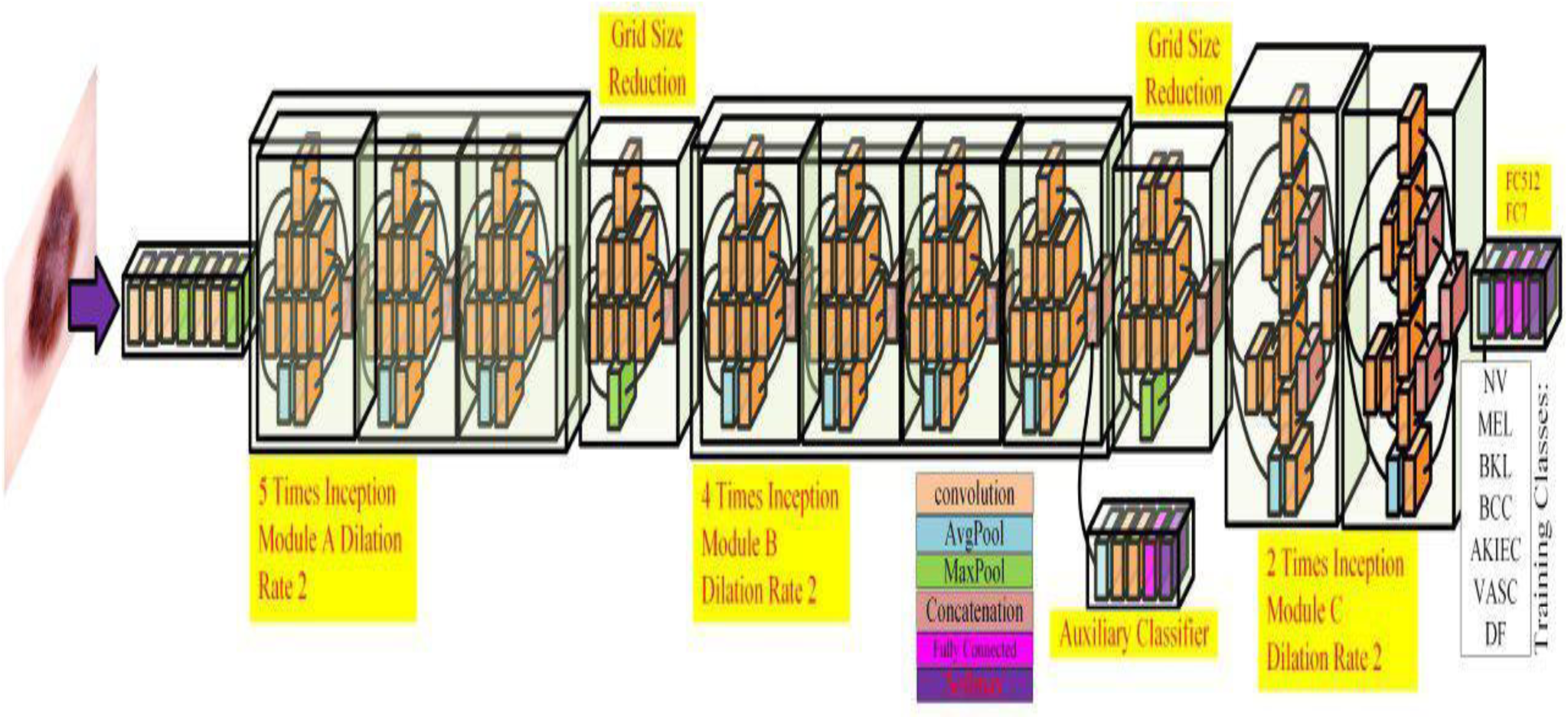
Dilated InceptionV3 network with three different modules of Inception blocks (5 times inception, 4 times Inception, 2 times inception). Every layer of Module A, Module B, and Module C has dilation rate (2,2)

**Fig. 7.**
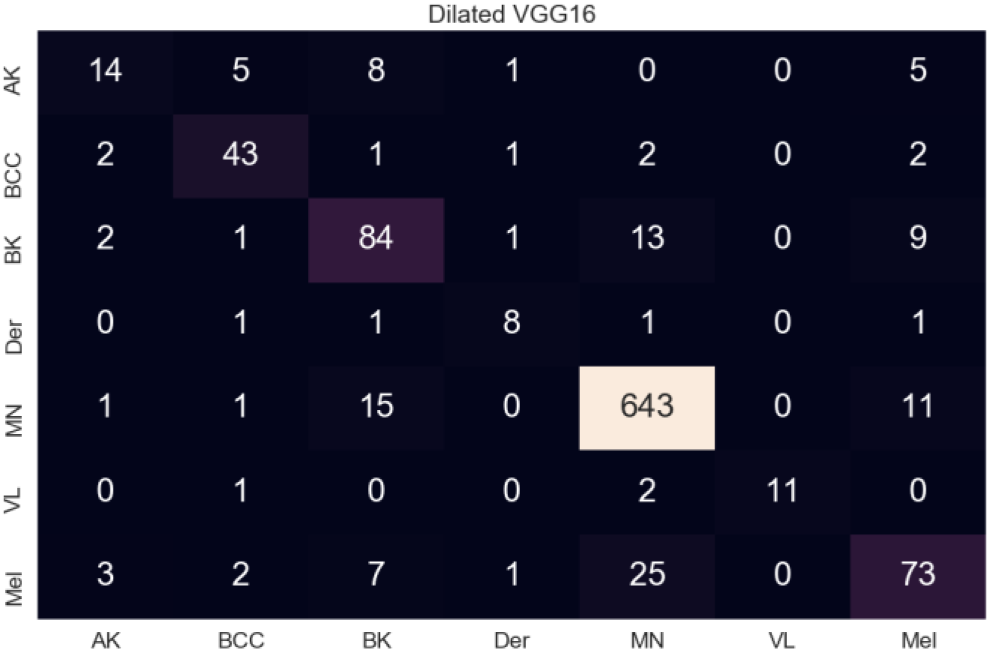
Confusion Matrix for proposed dilated VGG16

**Fig. 8.**
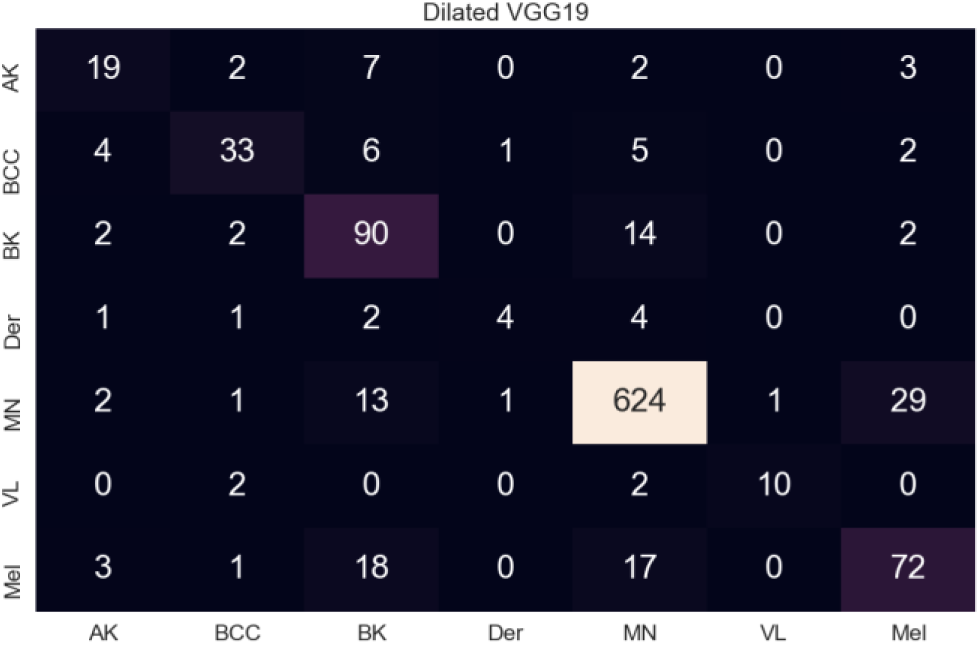
Confusion Matrix for proposed dilated VGG19

**Fig. 9.**
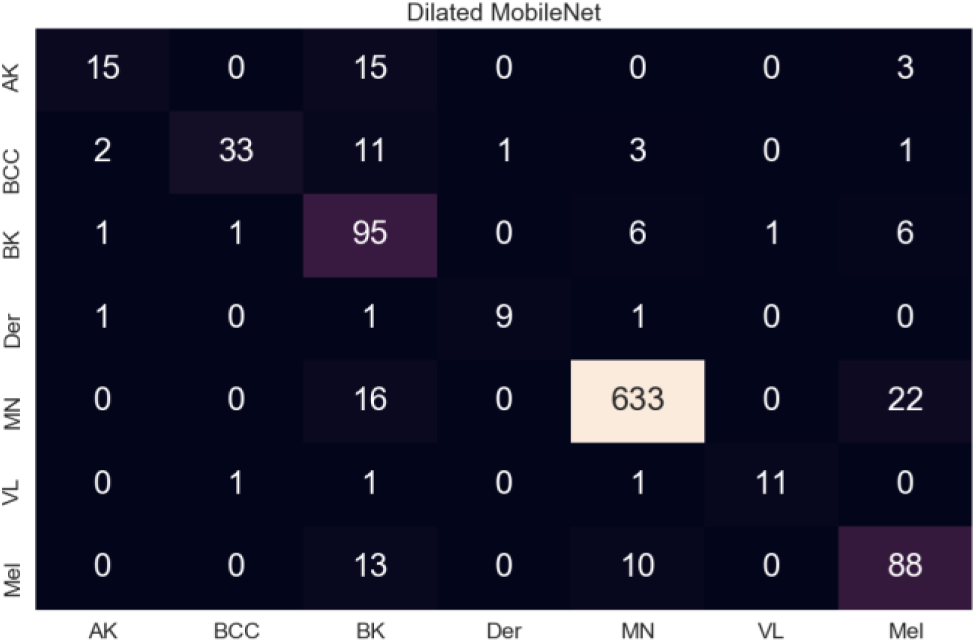
Confusion Matrix for proposed dilated MobileNet

**Fig. 10.**
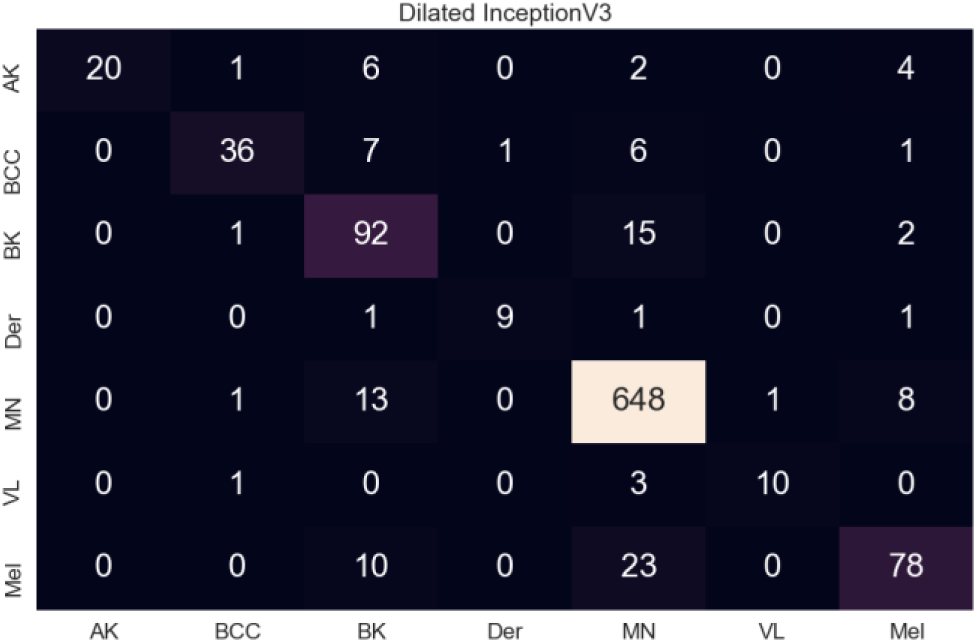
Confusion Matrix for proposed dilated InceptionV3

### D. Compare with Recent Work

Here, we compare our outcomes with some recent architectures on the HAM10000 dataset. For comparison, we use the recall, weighted average of precision, and the f-1 score of these seven different classes of the HAM10000 dataset. The proposed dilated InceptionV3 network produces superior outcomes in these three evaluation criteria than existing architectures. To compare with these state-of-art architectures, we evaluate average recall, average precision, and average F-1 score.

## VI. Conclusion and future work

All over the world, skin cancer is considered one of the deadliest types of cancer. Here, we construct a computer-aided skin lesion classifier system using four different dilated deep neural network architectures (VGG16, VGG19, MobileNet, and InceptionV3) with transfer learning techniques. Several different data preprocessing and augmentation rules applied to lessen the effect of class imbalance characteristic of HAM10000. We tried several evaluation approaches such as top-1 accuracy, recall, precision, f-1 score, and confusion matrix to compare our proposed model with the basic one. These models produce better outcomes than any known methods on skin lesions classification after considering image noise presence, the number of classes, and the issue of class imbalance. Among all the proposed architectures, InceptionV3 delivered superior classification accuracy, and MobileNet exhibits fewer parameters.

There are still many shortcomings exist which we will have to fix in the future. Every proposed architecture has been pre-trained with the ImageNet dataset; however, ImageNet is not well assembling for skin lesion images. Thus, utilizing a fine-tuning method to provide every layer an additional weight which increases the time and space complexities. Hence, to build a suitable application for medical, we need to shrink all the complexities.

